# The human placenta exhibits a unique transcriptomic void

**DOI:** 10.1101/2022.07.01.498408

**Authors:** Sungsam Gong, Francesca Gaccioli, Irving L.M.H. Aye, Giulia Avellino, Emma Cook, Andrew R.J. Lawson, Luke M.R. Harvey, Gordon C.S. Smith, D. Stephen Charnock-Jones

## Abstract

We have recently demonstrated that the human placenta exhibits a unique genomic architecture with an unexpectedly high mutation burden(Coorens et al. 2021) and it is also well recognized that the placenta uniquely expresses many genes(Gong et al. 2021). However, the placenta is relatively understudied in systematic comparisons of gene expression in different organs. The aim of the present study was to identify transcripts which were uniquely absent or depleted, comparing the placenta with 46 other human organs. Here we show that 40/46 of the other organs had no transcripts which were selectively depleted and that of the remaining six, the liver had the largest number with 26. In contrast, the term placenta had 762 depleted transcripts. Gene Ontology analysis of this depleted set highlighted multiple pathways reflecting known unique elements of placental physiology. However, analysis of term samples demonstrated massive over representation of genes involved in mitochondrial function (*P*=5.8×10^−10^), including PGC-1α - the master regulator of mitochondrial biogenesis, and genes involved in polyamine metabolism (*P*=2.1×10^−4^). We conclude that the term placenta exhibits a unique metabolic environment.

## Introduction

The placenta has a key role in the pathogenesis of many major complications of pregnancy, such as preeclampsia (PE) and fetal growth restriction (FGR), termed, collectively, the “Great Obstetrical Syndromes”(Brosens et al. 2011) and which account for a substantial burden of global morbidity and mortality. Progress on predicting and preventing these complications is hampered by lack of mechanistic understanding of normal and abnormal placental function and we and others have applied multiple studies using omic methods to try and address this knowledge gap. Published studies of the placenta transcriptome tend to focus on identifying genes differentially regulated in complicated pregnancies. Other studies have compared the placental transcriptome across species(Armstrong et al. 2017) and across gestation(Buckberry et al. 2017) but there are fewer studies comparing the placental transcriptome with the transcriptomes of other organs(Kim et al. 2012; Gong et al. 2021). RNA-Seq enables transcriptome profiling of tissues or single cells and there are a number of studies characterizing so-called the transcriptome ‘landscape’ of tissues of interest. It is now an essential part of large-scale multi-omics studies, such as the Encyclopedia of DNA Elements (ENCODE)(ENCODE Project Consortium 2012), the RoadMap Epigenomics Project(Roadmap Epigenomics Consortium et al. 2015), and the Functional Annotation of Mammalian Genome (FANTOM5) project(de Rie et al. 2017). However, the human placenta transcriptome is relatively understudied and absent from large-scale “omic” analyses such as the Genotype-Tissue Expression (GTEx) project(GTEx Consortium 2020).

Pan-tissue comparative analyses generally focus on identifying transcripts that are abundant in a tissue of interest while being absent or depleted in others. Indeed, there are a number of tools and databases that enable “tissue-specific” gene enrichment analysis(Jain and Tuteja 2019; Watanabe et al. 2019; Papatheodorou et al. 2020). Studying “tissue-specific” genes provides information about specific functions that define a unique set of characteristics or “identity” of a tissue of interest. Transcripts that are ubiquitously expressed in multiple tissues, such as house-keeping genes(Eisenberg and Levanon 2013) can be identified and this gives insight to the functions that all tissue share. In contrast, little attention has been paid to the identification of transcripts that are less abundant, or even absent, in one tissue compared to all others. Here we report transcripts depleted or absent in the human placenta at term and in early gestation compared with 46 other tissues studied in the GTEx project. Functional enrichment analysis of depleted transcripts highlighted pathways which reflect known aspects of placental physiology, such as lack of nervous tissue and unique immunological features. However, these analyses also generated evidence that the term human placenta has unique metabolic characteristics, as evidenced by multiple absent transcripts involved in mitochondrial function and polyamine metabolism.

## Results

### Tissue-wide comparison of depleted transcripts

We carried out mRNA sequencing (RNA-seq) using 59 human term placentas from the POP study cohort(Pasupathy et al. 2008; Gaccioli et al. 2016; Gong et al. 2018b) and 14 human placentas from earlier in gestation (n=8, 7-8 weeks (8wk); n=6, 13-14 weeks (14wk))(Prater et al. 2021). We obtained approximately 38 million reads from each sample (**Supplementary Table 1**). We compared the placental transcriptome profile at 8wk, 14wk and term with that of 46 tissues from 11,803 samples of GTEx Consortium datasets(GTEx Consortium 2020) and investigated which transcripts are absent or depleted in the placenta while being reasonably abundant in other tissues (**Supplementary Table 2**). To adjust for differences in the RNA composition across tissues, we applied the following two normalization methods: 1) the median ratio method (DESeq(Anders and Huber 2010)) and 2) the trimmed mean of M-values (TMM(Robinson and Oshlack 2010)) (see Methods for details). For 19,170 eligible protein-coding transcripts, we ranked tissues by their normalized count per million (nCPM) and identified 5,632 and 5,727 transcripts for which the term placenta was ranked 47 (i.e. bottom) based on DESeq and TMM normalization methods, respectively. Then we selected the 762 transcripts which satisfied the following three conditions: (1) nCPM >1 for the tissue of rank 46, (2) nCPM_(rank=46)_/nCPM_(rank=47)_ > 3, and (3) fulfilling these conditions both in DESeq and TMM normalization methods (**Supplementary Table 3**) – these transcripts are described as being “depleted” hereafter. For early gestational age datasets (8wk and 14wk placenta) we identified 215 and 112 such depleted transcripts respectively (**Supplementary Tables 4 and 5**).

Using the same criteria we applied to the placenta, we sought to identify mRNAs depleted in each of the 46 other tissues. Surprisingly, we found that there were ∼30 times more transcripts depleted in the placenta than in the liver (26 depleted transcripts), which was the highest among the 46 non-placental tissues. Besides the placenta, only six tissues had one or more depleted transcript: liver, spleen, brain (cerebellar hemisphere), muscle, testis and adrenal gland (**Figure 1A** and **Supplementary Table 6**). We then used five external placenta RNA-Seq datasets generated independently (two early gestational placenta datasets: (1) Lim et al.(Lim et al. 2017) (n=4) and (2) Huang et al.(Huang et al. 2018) (n=3) and three term placenta datasets: (3) Verheecke et al.(Verheecke et al. 2018) (n=66), (4) Ashley et al.(Ashley et al. 2021) (n=4), and (5) Awamleh et al.(Awamleh et al. 2019) (n= 21)). These analyses confirmed that the placenta had the most depleted transcripts among the other tissues studied −132 (Lim), 55 (Huang), 237 (Verheecke), 340 (Ashley), and 123 (Awamleh) (**Figure 1A** and **Supplementary Table 7**). All the datasets described above had variable depth of coverage. So, to investigate any possible effect of sequencing depth, we down-sampled the reads to 20 million for all samples. We then repeated the analysis with the same methods and criteria as described above. In down-sampled datasets, we obtained consistent results that the placenta had many more depleted transcripts than other tissues (**Supplementary Figure 1** and **Supplementary Table 8**). The number of depleted transcripts that were shared among our dataset and other term placenta datasets (**Figure 1B**) or other first trimester datasets (**Figure 1C**) was highly significant (*P*<1×10^−314^ for the term placenta and *P*=7.4×10^−209^ for the early placenta; both Fisher’s exact test). In our term and early gestation placental datasets, there was a total of 883 transcripts depleted at any gestational age and among these, 58 were depleted in all three trimesters (**Supplementary Table 9** and **Figure 1D**). **Figure 1E** shows the 58 genes for which the transcripts are depleted in our three placental datasets ranked by their fold change, with the top 10 genes being annotated, compared to 46 tissues from the GTEx dataset (see **Supplementary Table 9**).

**Figure 1.**
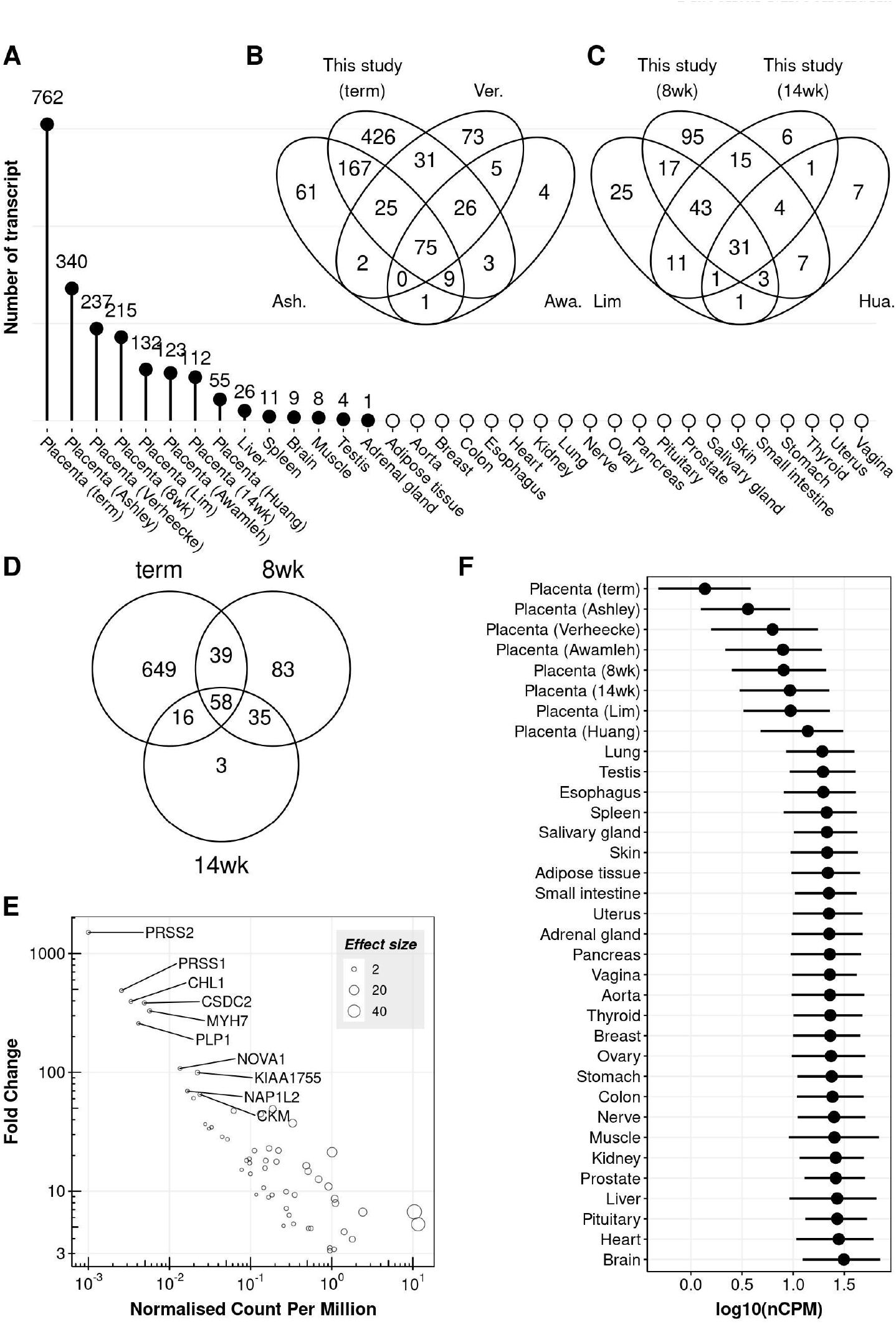
The number of absent or depleted transcripts in various tissues. **A**, The number of depleted transcripts are shown in the placenta samples (our and other studies) and other somatic tissues. The term, 8wk and 14wk placentas are f rom this study. Tissues with open circles represent zero depleted transcripts. **B**-**D**, Venn diagrams showing the number of transcripts, and their overlaps, depleted in early gestational placentas (**B**), term placentas (**C**) and three gestational stages of the placenta datasets f rom this study (**D**). The lists of transcripts depleted in the non-placental and the external placental tissues are available in **Supplementary Tables 6 and 7**, respectively. Ver. (Verheecke), Ash. (Ashley), Awa. (Awamleh), Hua. (Huang). **E**, Abundances of the transcripts (x-axis) relatively depleted in all three trimesters are shown along with their fold change (y-axis, calculated as follows: nCPM_(rank=46)_/nCPM_(rank=47; placenta)_). The counts (per million) on x-axis are normalized by TMM method and the data is available in **Supplementary Table 9**. To avoid fold change being infinite values, a small number (0.001) was added to nCPM of the term placenta. The transcripts with the 10 highest fold change (*PRSS2* being the top one) are shown with their gene names. The size of circle represents the effect size (i.e. nCPM_(rank=46)_ -nCPM_(rank=47; placenta)_). **F**, The range of transcript abundances for the 762 genes in the placental tissues and 26 representative somatic tissues out of 46 we studied. Dot: median; line: interquartile range (IQR). For display, non-placental tissues shown in **F**, were manually selected if there are at least two subregions f rom the same tissue. For example, we analyzed a total of 13 brain subregions in this study and the cerebellar hemisphere is shown here to represent the brain. The representative sub-regions are shown in **Supplementary Table 2**.

Among the 762 transcripts depleted in our term placenta dataset, we did not detect any transcripts encoding *PRSS2* (serine protease 2 also known as Trypsin 2) whilst *MT-ND6* (mitochondrially encoded NADH:ubiquinone oxidoreductase core subunit 6) had the highest level of expression (nCPM=123), but was still three-fold lower than in all the other tissues studied (**Supplementary Table 3**). Interestingly, the 762 transcripts depleted in our term placenta were also less abundant in our early gestation placenta samples and other placenta data sets studied (**Figure 1F**). Moreover, the 426 transcripts uniquely depleted in our term placenta data (**Figure 1B**) were also less abundant in external term placenta datasets and majority of them were ranked bottom in the Ashely (335; 79%), Verheecke (245; 58%), and Awamleh (179; 42%) datasets (**Supplementary Figure 2**).

### Dynamic changes of depleted transcripts during pregnancy

We further investigated the 883 transcripts depleted in the placenta at any trimester of pregnancy (**Figure 1D**) to see how they change over time. Among the 215 transcripts depleted in the first trimester placenta, 83 transcripts were not shared with other gestational ages (T1 in **Figure 2A**) and their abundances increased as gestation progressed (**Supplementary Figure 3A**). A number of gene ontology (GO) terms were significantly enriched (*P*<0.05) in these 83 transcripts (T1 in **Figure 2B**), including MHC protein complex (*HLA-A, HLA-DPA1* and *HLA-DPB1*) and ion channel inhibitor activity (*ANKRD36, CAMK2D, LYNX1*, and *SCN1B*), suggesting a limited role for these functions in the first trimester.

**Figure 2.**
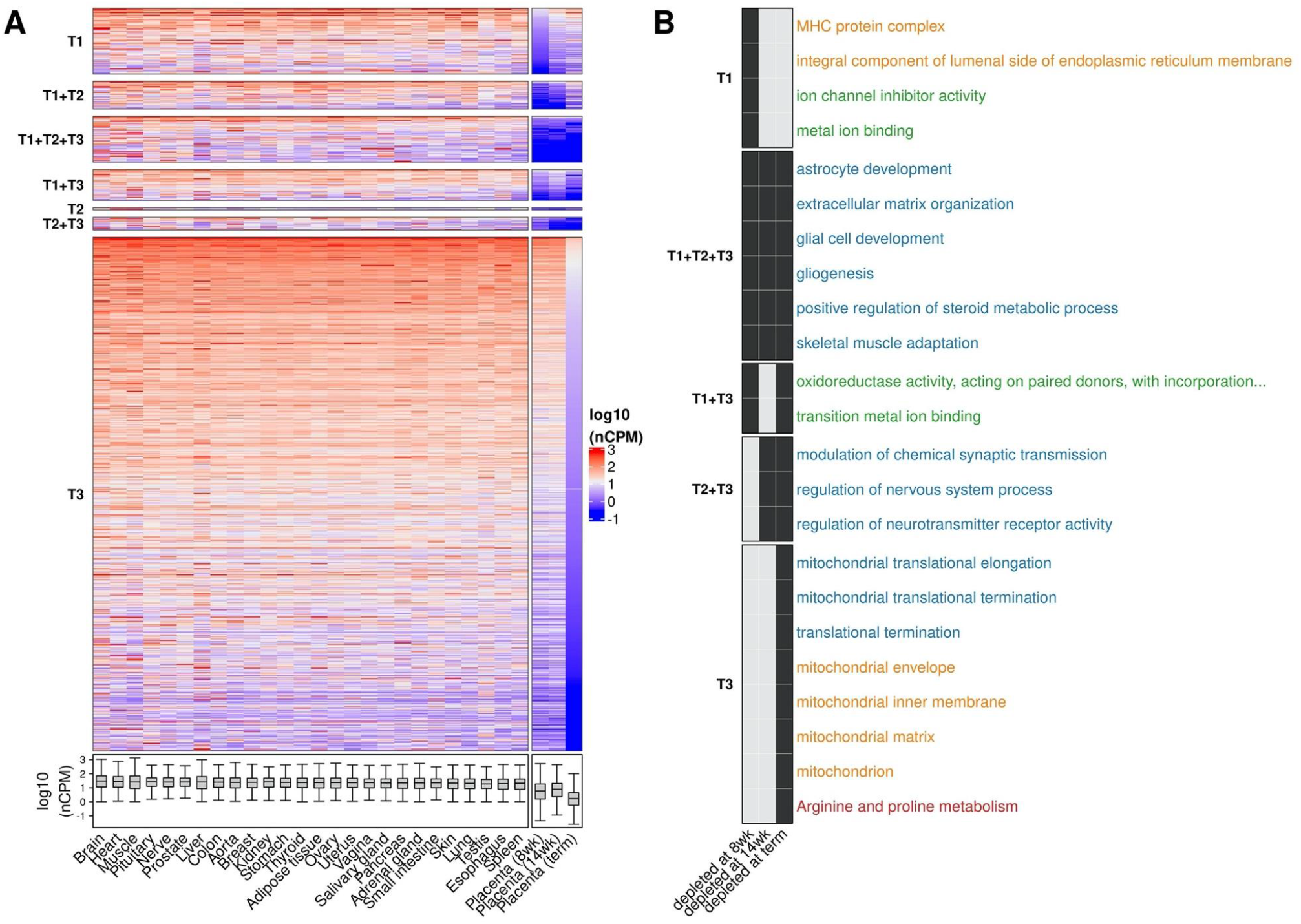
Abundance and GO analysis of transcripts depleted in the placenta during pregnancy. **A**, A heatmap representation of the abundance of 883 placenta-depleted transcripts (rows) across 29 tissues including our placenta datasets, i.e. 8wk (T1), 14wk (T2), and term (T3). nCPM (log_10_ scale) is color-coded f rom red (higher) to blue (lower) and their values across the 883 transcripts are shown as a boxplot in the bottom. 26 non-placental tissues are shown in descending order of the median CPM and the placenta samples are ordered by gestational age. The 883 transcripts are ordered in descending order of the average CPM across the placental samples. **B**, A summary of the gene ontology (GO) and KEGG pathway analysis. Significantly (adjusted *P*<0.05) overrepresented GO terms and KEGG pathway are shown with the following three sources: biological process (blue); molecular function (green); cellular component (orange); KEGG (red). The black and grey square represent being depleted and not being depleted, respectively, at the corresponding gestational age. To note, GO terms with at least the depth of 5 or more f rom the root and the intersection size of 3 or more (i.e. the number of overlaps between the query and the number of annotated genes belong to the GO terms in question) were plotted. For KEGG pathways, those with the intersection size of 5 or more were plotted. The full list of significantly overrepresented GO terms is shown in **Supplementary Table 10**. For **A** and **B**, the 883 transcripts are divided by their depleted status among the placenta samples at 8wk, 14wk, and term as shown by the following: T1: depleted at 8wk only; T1+T2: depleted at 8wk and 14wk; T1+T2+T3: depleted at 8wk, 14wk and term; T1+T3: depleted at 8wk and term; T2: depleted at 14wk only; T2+T3: depleted at 14wk and term; T3: depleted at term only. For display, non-placental tissues, shown in **A**, were manually selected if there are at least two subregions f rom the same tissue. For example, we analyzed a total of 13 brain subregions in this study and the cerebellar hemisphere is shown here to represent the brain. The representative sub-regions are show in **Supplementary Table 2**. nCPM: normalized count per million; wk: week of gestation.

We found the second trimester placenta sample had the smallest number of depleted transcripts (n=112) compared to those from the first trimester (n=215) and the term placenta (n=762). Indeed only 3 transcripts (cilia and flagella associated protein 91 (*CFAP91*, also known as *MAATS1*), myomesin 2 (*MYOM2*), and neurotrophic receptor tyrosine kinase 3 (*NTRK3*)) were uniquely depleted in the second trimester (T2 in **Figure 2A**). However, they all were ranked bottom (i.e. the least abundant compared to non-placental tissues) in the first trimester and close to bottom in the term placenta. However, they were not sufficiently low enough to satisfy the 3-fold threshold for both TMM and DESeq normalization methods. For example, for *MYOM2* the fold changes based on TMM and DESeq were 2.9 and 2.8 fold respectively.

Among the 883 transcripts depleted in any of the three trimesters, 58 are depleted in all three (**Supplementary Table 9** and T1+T2+T3 in **Figure 2A**) and they are associated with various GO terms (T1+T2+T3 in **Figure 2B**). It is unsurprising that genes annotated with the GO terms “astrocyte development, “gliogenesis” and “skeletal muscle adaptation” are absent in the placenta. However, as the placenta is a steroidogenic organ the depletion of genes associated with “positive regulation of steroid hormone metabolic process” is more surprising. The genes annotated with this term include aldo-keto reductase family members (*AKR1C1* and *AKR1C2*). This is consistent with the requirement for placental steroid production as these reductases inactivate steroid hormones(Penning et al. 2015), specifically progesterone in the case of *AKR1C1*. Peroxisome Proliferator Activated Receptor Gamma (PPARG) is abundant and is essential for placental development(Valle et al. 2005) and function. However, the depletion of PPARG Coactivator 1 Alpha (*PPARGC1A)* transcripts (also known as PGC-1α) suggests that the usual coordination between PPARG and PGC-1α does not occur in the placenta(Hondares et al. 2006). Of note, PGC-1α is also directly implicated in regulating mitochondrial biogenesis(Wu et al. 1999) and the regulation of mitochondrial genes (see below).

Several depleted transcripts are annotated with the GO term “extracellular matrix organization”. Trypsins 1 and 2 (PRSS1 and PRSS2) are notable as these were essentially absent from the placenta. As these proteins are key activators of multiple matrix metalloproteases this suggests that initiation of matrix remodeling is mediated by other proteases. Transcripts encoding two type IV collagen genes (*COL4A3, COL4A4*) were depleted. These collagens are components of basement membranes and form a triple helix (with COL4A5). Mutation or loss of any of these three genes causes Alport’s syndrome(Hudson et al. 2003). However, the lack of the α3. α4. α5(IV) collagen protomer is without effect in the placenta, in contrast to the other organs affected in Alport’s syndrome. It is likely that the α1. α1. α2(IV) collagen protomer is sufficient and in fact the placenta has the highest expression of COL4A1 among the GTEx tissues from our previous study(Gong et al. 2021). Keratin filament transcripts (*KRT4, KRT5, KRT13*) are also depleted and annotated with the GO term “extracellular matrix organization”. These keratins are characteristic of stratified epithelial surfaces(Moll et al. 2008) (such as the esophagus in which the expression level is >10,000 times higher) and this difference likely reflects the syncytial nature of the trophoblast epithelial surface.

We identified 762 depleted transcripts in the term placenta and this was the highest number among three trimesters, and 649 of them (85%) were uniquely depleted at term (T3 in **Figure 2A**). GO analysis showed these genes are predominantly associated with mitochondria-specific processes, suggesting that the term placenta has diminished capacity for these functions (T3 in **Figure 2B**). They include genes encoding 12 mitochondrial ribosomal proteins, ATP synthase, H^+^ transporting, mitochondrial F1 complex, delta subunit (*ATP5D*), succinate dehydrogenase complex assembly factor 1 (*SDHAF1*), NADH:ubiquinone oxidoreductase complex assembly factors (*NDUFAF3, NDUFAF8*) and subunits (*NDUFB7, NDUFS7, NDUFS8*), and mitochondrially encoded NADH:ubiquinone oxidoreductase core subunit 6 (*MT-ND6*). Even though these transcripts were not sufficiently low to be classified as being depleted in earlier gestations, most of them were also less abundant than somatic tissues (**Supplementary Figure 3B**). Using KEGG (Kyoto Encyclopedia of Genes and Genomes) pathway analysis of the 649 transcripts depleted only at term, we also noted that genes for arginine and proline metabolism and hence the polyamine (putrescine, spermidine, and spermine) metabolic pathway were also significantly over-represented (**Supplementary Text and Supplementary Figure 4**).

Having observed significantly over-represented mitochondria-related GO terms in the list of depleted transcripts, we examined the proportion of RNA-Seq reads from the 19,170 protein-coding genes that mapped to mitochondrial DNA. The term placenta has the lowest percentage (3.4%), followed by the aorta (4.5%), and the 8wk placenta (4.6%) (**Supplementary Figure 5A**). In contrast, the heart (left ventricle; 39.7%), the kidney (cortex; 31.3%) and the liver (21.1%) expressed the most mitochondrial protein-coding transcripts. We also examined the extent of mitochondrial transcripts including both the protein-coding and non-coding transcripts, such as mitochondrial rRNA and tRNA, and found the term placenta also showed the lowest proportion of reads mapped to mitochondrial DNA (3.7%, **Supplementary Figure 5B**).

### Abundance of nuclear-encoded transcripts localized in the mitochondria

Mitochondria contain proteins encoded by nuclear DNA and subsequently imported to the mitochondria as well as those directly encoded by mitochondrial DNA (mtDNA). MitoMiner(Smith and Robinson 2016) is a database of protein coding genes with strong support for mitochondrial localization and hence function. Having observed association of mitochondria-related GO terms in the 762 transcripts depleted in the term placenta, we further investigated how many of these encode mitochondrial proteins defined in MitoMiner and found 84 (*P*=5.1×10^−10^, Fisher’s exact test). For the 234 transcripts depleted earlier in gestation (either first or second trimester, **Figure 1D**), the number overlapping with MitoMiner is only 9 (*P*= 0.382, Fisher’s exact test). We then examined the abundance of transcripts encoding the 1,042 genes in MitoMiner across non-placental tissues from GTEx and placentas obtained in all three trimesters (8wk, 14wk and at term). We found that the 3 placental tissues clustered on a single branch and were distinct from the other tissues (**Figure 3A**). The transcript abundance of the 1,042 MitoMiner genes were lowest in the term placenta, followed by the 14wk placenta, while they were the most abundant in the muscle, followed by the liver and the heart (**Supplementary Figure 6)**. However, the term placenta also had 9 transcripts whose abundance levels were higher than any other somatic tissues we compared (**Figure 3B**) – two of them (KMO and ARMS2) were measured using RT-qPCR (discussed further below). Interestingly multidimensional scaling of all 11,876 samples showed a profound clustering of all 49 tissues, indicating tissue-specific expression of the 1,042 genes (**Figure 3C** and **Supplementary Figure 7**). All placental samples were clustered closely together. Next, using whole-genome sequencing (WGS) datasets, we examined mtDNA copy numbers of the term placental tissue (n=80) and compared those of four healthy tissues (endometrium(Moore et al. 2020) (n=398), blood(Lee-Six et al. 2018) (n=199), colon(Lee-Six et al. 2019) (n=568), and liver(Brunner et al. 2019) (n=517); see **Supplementary Table 11, Supplementary Figure 8**) and 21 non-placental tissues from the Cancer Genome Atlas Pan-Cancer Analysis of Whole Genomes (PCAWG) Consortium(Yuan et al. 2020). We found that mtDNA copy number was not substantially lower in the placenta than other tissues we compared (see **Supplementary Text**).

**Figure 3.**
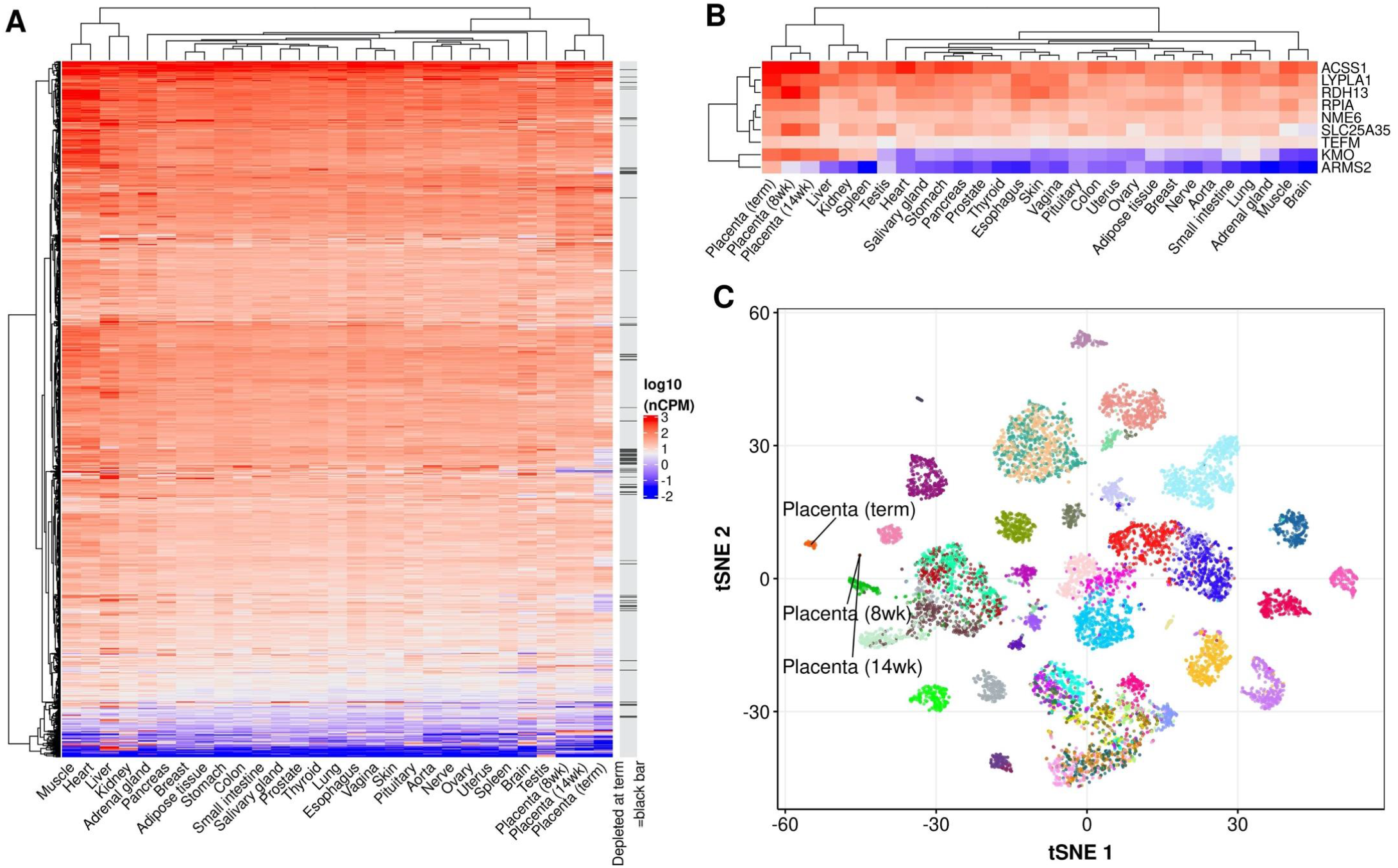
Nuclear-encoded mitochondrial transcripts. **A**, A heatmap representation of the abundance of 1,042 genes in MitoMiner (rows) across 29 tissues (columns). nCPM (log _10_ scale) of a transcript is color-coded f rom red (higher) to blue (lower). **B**, A heatmap showing the abundance (log _10_ scale) of 9 highly enriched MitoMiner transcripts in the placenta. **C**, A multidimensional scaling plot (tSNE) of 11,876 samples f rom 49 tissues using nCPMs (calculated by TMM method) of 1,042 genes in MitoMiner. Each dot represents one of 11,876 samples (i.e. 11,803 samples f rom GTEx and 73 f rom the placenta of the following gestational ages: 8 at 8wk, 6 at 14wk and 59 at term) and each color represents a tissue and the full color-coding is shown in **Supplementary Figure 7**. For display, non-placental tissues, shown in **A**, were manually selected if there are at least two subregions f rom the same tissue. For example, we analyzed a total of 13 brain subregions in this study and the cerebellar hemisphere is shown here to represent the brain. The representative sub -regions are show in **Supplementary Table 2**. nCPM: normalized count per million; wk: week of gestation; tSNE: t - distributed stochastic neighbor embedding.

### Validation of placental transcript abundances

To confirm the transcript abundance levels of the placenta and other somatic tissues that we identified based on RNA-Seq dataset, we performed RT-qPCR assays in independent samples. We selected a total of 13 transcripts (11 depleted and 2 mitochondria-associated transcripts enriched in the term placenta) and measured their mRNA levels in eight human tissues: placenta (term), aorta, heart, breast, lung, stomach, small intestine and colon (see Methods for details). For each of the 11 depleted transcripts, we confirmed that their abundances were significantly lower in the placenta than the other tissues (*P*=1.7×10^−5^, Mann-Whitney test) and the placenta was ranked lowest (**Figure 4**). We also confirmed that the mRNA level of KMO, one of the enriched transcripts in the placenta, was the highest in the placenta and it was significantly higher (*P*=2×10^−5^, Mann-Whitney test) than the other tissues tested (**Figure 4**). For ARMS2, another enriched target, 19 of the 24 non-placental tissue samples used were not assayable by qPCR, (below the limit of detection), whereas all 5 of the placental samples were measurable – this is consistent with the RNA-Seq data showing enrichment in the placenta (**Figure 3B**). Overall, the 13 transcripts selected for validation of either being depleted (11) or enriched (2) from RNA-Seq datasets were confirmed using RT-qPCR.

**Figure 4.**
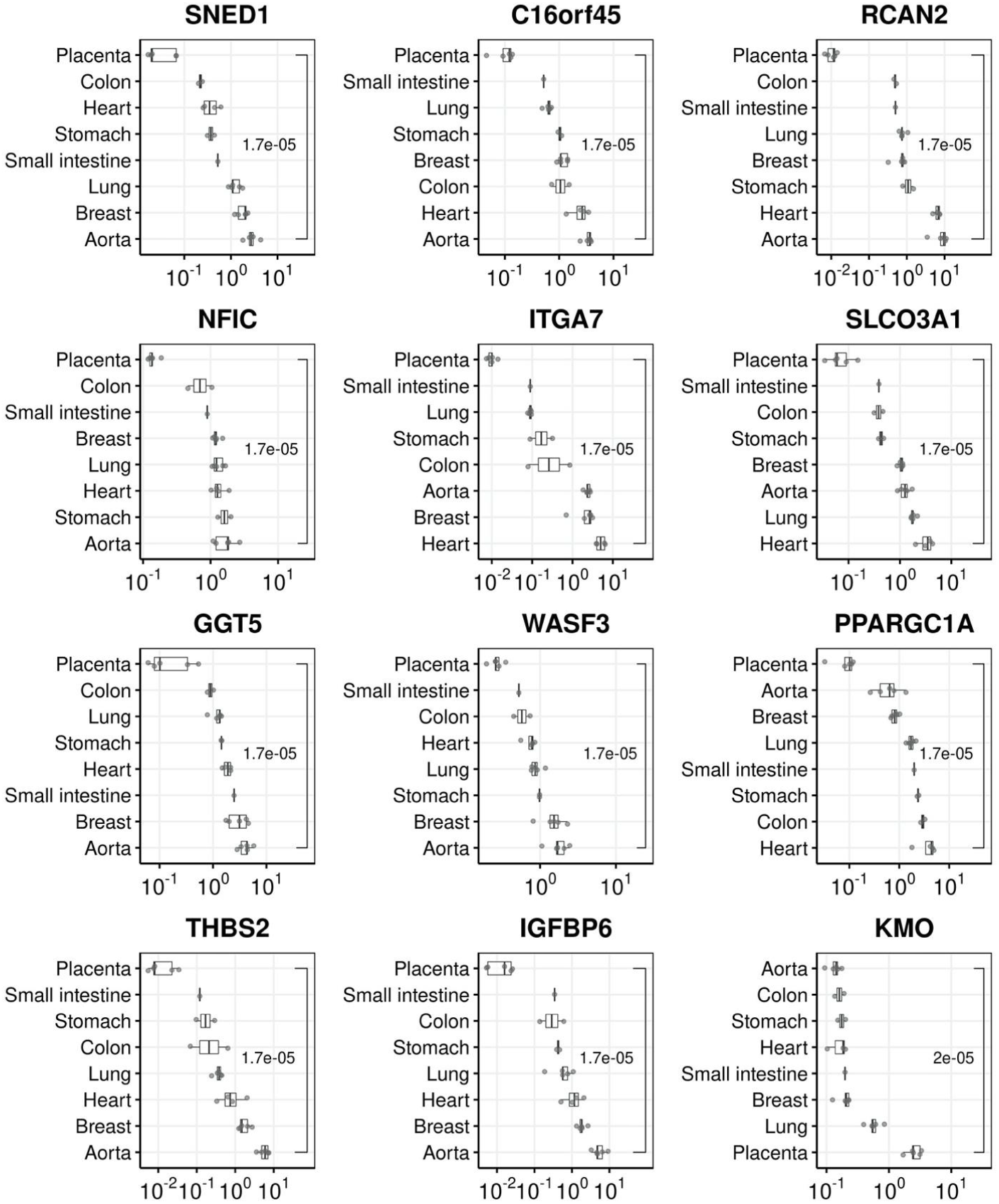
qPCR validation assays for 14 transcripts. The transcript abundance levels (x-axis) were measured in 8 tissues (y-axis) which were ranked f rom the lowest to highest level using the median abundance for each transcript. Each dot represents an individual sample (by taking the mean of the technical triplicate) and each of the boxes shows the median and interquartile range (IQR). The horizontal lines (whiskers) extended f rom the box represent a range of 1.5*IQR f rom both ends. The P - values (Mann-Whitney test) are shown between the placenta and the rest of non-placental tissues. The qPCR data used for the graph are available in **Supplementary Table 12**.

## Discussion

The key finding of the present study is that the human placenta has a unique transcriptome architecture compared to all of the organs studied in the GTEx project. More than 80% of the organs studied in the GTEx project lack even a single uniquely depleted transcript. Of the organs studied in the GTEx project, the liver had the largest number of uniquely depleted transcripts, with 26 depleted or absent. Strikingly, the transcriptome of term placenta had almost 30 times more depleted or absent transcripts than the liver. Gene Ontology analysis indicated that some of the absent transcripts reflect known unique qualities of placental function. For example, the placenta has no innervation and this likely reflects the identification by Gene Ontology analysis of multiple transcripts involved in formation of elements of the nervous system. Similarly, recognition of the allo-immune placenta is essential for normal pregnancy(Moffett et al. 2017) and this is reflected in unique expression of MHC antigens, which was another Gene Ontology analysis pathway identified. However, Gene Ontology analysis identified other pathways which we did not anticipate.

The most striking example was that many transcripts related to mitochondrial function were far less abundant in the term placenta than other somatic tissues. Interestingly, the mRNA for PPARGC1A (Peroxisome proliferator-activated receptor gamma coactivator 1-alpha; also known as PGC-1α), a transcriptional master regulator of mitochondrial biogenesis, was depleted in the placenta in all three trimesters, as well as in four external placenta datasets (**Supplementary Table 7**). *Ppargc1a*-null mice have decreased expression of mitochondrial genes, especially those encoding various subunits of the electron transport chain (Austin and St-Pierre 2012; Vernier and Giguère 2021), suggesting a possible link between its low abundance and the diminished content of mitochondrial transcripts in the placenta. PGC-1α interacts with a very wide range of transcriptional co-activators and is a key regulator of metabolic homeostasis(Miller et al. 2019) and protects cells against oxidative damage by inducing the expression of several ROS (reactive oxygen species) detoxifying enzymes such as superoxide dismutase 2 (SOD2). Interestingly, our RNA-Seq analysis showed that SOD2 mRNA was the lowest in the early gestation placentas as well as having a low rank (45/47) in the term placenta. As ectopic expression of PGC-1α reduced levels of ROS(Valle et al. 2005; St-Pierre et al. 2006), it has been suggested that PGC-1α ensures high energy metabolism and removal of its toxic by-products at the same time. We recently reported that the placenta has a unique somatic mutation profile(Coorens et al. 2021), predominantly the SBS18 signature which is associated with oxidative stress. This could be explained, at least in part, by accumulation of ROS possibly due to lower level of PGC-1α and SOD2 transcripts in the placenta.

The syncytiotrophoblast (STB) is a multinucleated epithelium covering the outer layer of chorionic villi and it differentiates from cytotrophoblast (CTB). The STB mitochondria have different morphological properties compared to CTB, specifically, they are smaller with irregular spherical cristae and a dense matrix(Holland et al. 2017; Fisher et al. 2020) and it has been suggested that these changes are related to steroidogenesis of STB(Martínez et al. 1997; Martinez et al. 2015). Functional studies have shown that STB mitochondria have reduced membrane potential, increased levels of hydrogen peroxide, lower antioxidant level and are more sensitive to ROS(Watson et al. 1998; Bustamante et al. 2014; Schoots et al. 2018). These characteristics are due to the complement of mitochondrial proteins present and again reflect the features of placental biology that are not found in other tissues.

We have previously reported that placental polyamine metabolism is implicated in placentally related complications of human pregnancy(Gong et al. 2018b). In the current study we found that genes associated with the KEGG pathway “arginine and proline metabolism” were over-represented among the depleted transcripts. Within this group were transcripts from five genes (SRM, SAT2, SMOX, AZIN2, and PAOX) involved in polyamine metabolism; these were depleted at term but were also less abundant earlier in gestation (see **Supplementary Text** and **Supplementary Figure 4**). Interestingly, expression levels of some genes in the polyamine pathway (e.g., *SAT1, AZIN1, SMS*, and *AMD1*) were higher in the placenta than non-placental tissues. Kajander et al.(Kajander et al. 1989) reported enzymatic activities of spermidine synthase (SRM) and spermine synthase (SMS) in seven human tissues, and the SMS-to-SRM ratio was the highest in the placenta (∼5) followed by the kidney (∼3.7). In our comparative analysis of RNA-Seq datasets, we confirmed that the SMS-to-SRM ratio at their transcript level was also the highest at the term placenta (48.7), followed by the 14wk placenta (5.4), and the 8wk placenta (5.3). The kidney cortex (2.5) appeared to have the highest ratio among 46 non-placenta tissues we compared, with the pancreas (0.17) being the lowest – it was also the lowest from the protein ratio (0.4) reported by Kajander. This demonstrates that our tissue-wide comparison of transcripts parallels previous analyses based on measurement of proteins level.

The mechanisms underlying the reduced expression of the selectively depleted transcripts in the placenta remains to be determined. One we considered was differential methylation of the promoter regions of such genes. While it has long been recognized that placental DNA is globally hypomethylated compared to other tissues(Ehrlich et al. 1982; Fuke et al. 2004), it varies in locus-specific manner(Chatterjee et al. 2016; Gong et al. 2018a). We previously reported 71 protein-coding transcripts specifically enriched in the placenta(Gong et al. 2021), and we compared their promotor methylation levels(Gong et al. 2018a) with those of the transcripts depleted in the placenta we report here. However, we did not observe any significant difference in DNA methylation between the enriched and depleted transcripts in the CPG islands associated with these genes (**Supplementary Figure 9**). Moreover, when we studied the promoter regions of the two sets of genes, the median promoter methylation of enriched transcripts (36.9%) was actually higher than that of depleted transcripts (10.2%, *P*=3.4×10^−11^). This observation runs counter to the typical “high-methylation - low-expression” relationship and suggests that other mechanisms may be involved, and this is an area for future study. We considered possible differences in the number of mitochondria per nucleus but the placenta was not an outlier in mitochondrial DNA copy number (**Supplementary Figure 8**) and some mitochondrial genes actually had higher levels of expression in the placenta than in other tissues.

We noted that the proportion of genes encoding depleted transcripts from chromosome 19 was higher than expected by chance (Odds Ratio=3.18, *P*=1.5×10^−25^; Fisher’s exact test) (**Supplementary Figure 10**). This chromosome has the highest gene density of all human chromosomes and the highest GC and CpG content. It is unusual in that nearly a quarter of its genes fall in 20 tandemly arranged gene families(Grimwood et al. 2004; Harris et al. 2020). Several of these families have direct roles in pregnancy – for example, the beta subunit of chorionic gonadotropin (CGB, 6 functional genes), the pregnancy specific glycoproteins (PSG, 10 functional genes) and the large imprinted cluster of miRNAs which are largely placenta specific(Donker et al. 2012) (C19CM, 46 genes). These genes are all highly expressed in the placenta(Gong et al. 2021). However, chromosome 19 encodes other unusual gene families. Natural killer cells play an important role in human pregnancy(Colucci 2019) and their receptors (killer cell immunoglobulin-like receptors, KIRs.) are highly polymorphic and are in a cluster within the 1 Mb leukocyte receptor complex in Chromosome 19.

Finally, transposable elements or their remains account for a surprisingly large fraction of the human genome and host organisms have evolved numerous strategies to defend themselves against the threat posed by functional endogenous retroelements. One such mechanism involves the very large and rapidly evolving family of transcription factors, the Krüppel-associated box domain zinc finger proteins (KRAB-ZFPs)(Bruno et al. 2019). There are an estimated 352 genes encoding KRAB-ZFPs in the human genome, and 209 of these are located within six clusters on chromosome 19(Yang et al. 2017). Thus chromosome 19 has many unusual features and many of these are related to placental or reproductive function. We now add to its list of unusual features – that of having an over representation of depleted transcripts in the human placenta.

Clearly, there is a risk that apparent differences in expression could be observed due to batch effects and this could occur at any of a range of levels, including RNA extraction, the sequencing platform employed, the transcriptome size, and the normalization method employed. For example, the results of long RNA-seq are profoundly affected by whether RNA was size selected or selected by oligo dTs for selective extraction of mRNA through its polyadenylated tail(Gong et al. 2021). We mitigated these risks using multiple approaches. First, we ensured comparable methods of RNA extraction and sequencing between our own samples (mirVana miRNA isolation Kit, Ambion) and GTEx (Tissue miRNA Kit, PreAnalytix^®^ and miRNeasy Mini Kit, Qiagen). Second, we analyzed multiple distinct placental RNA-seq datasets, including those we generated and using other publicly available sources. Third, we compared multiple bioinformatic approaches to determine that the results were robust to definitions using different pipelines. Finally, we externally validated some of the key results using qPCR and a separate group of placental samples and tissue samples obtained from a local tissue bank. We conclude that these data support a unique transcriptomic void in the human placenta. We speculate that this void might identify transcripts that are dispensable in a transient organ – but not in others. Moreover, the neglect of the placenta in a large-scale international consortium resulted in the exclusion of one of the human body’s most interesting transcriptional landscapes.

## Materials and Methods

### Placenta samples

All the full-term placenta samples were obtained from the Pregnancy Outcome Prediction (POP) study, a prospective cohort study of nulliparous women attending the Rosie Hospital, Cambridge (UK) for their dating ultrasound scan between January 14, 2008, and July 31, 2012. The study has been previously described in detail(Pasupathy et al. 2008; Gaccioli et al. 2016). Ethical approval for the study was given by the Cambridgeshire 2 Research Ethics Committee (reference number 07/H0308/163) and all participants provided written informed consent. A total of 128 unique placental samples were analysed in this study - 60 samples were used for RNA-Seq (**Supplementary Table 1**) and 80 for WGS (**Supplementary Table 11**). One of the RNA-Seq samples was dropped from the analysis due to the presence of decidual contamination(Gong et al. 2021) and 12 placentas were analyzed using both methods.

First and second trimester tissue samples were collected with informed written patient consent and approval of the Joint University College London/University College London Hospital Committees on the Ethics of Human Research (05/Q0505/82) from 7-8 wGA (n=8) and 13-14 wGA (n=6) uncomplicated pregnancies. Gestational age was confirmed by ultrasound measurement of the crown-rump length of the embryo. All samples were collected from patients undergoing surgical pregnancy termination under general anesthesia for psycho-social reasons. Villous samples were obtained under transabdominal ultrasound guidance from the central region of the placenta using a chorionic villus sampling (CVS) technique. All samples were snap-frozen immediately in liquid nitrogen and stored at −80°C until analysis. These samples have previously been described in full(Prater et al. 2021).

### RNA sequencing and data processing

The POP study placental biopsies were collected within 30 minutes of delivery and flash frozen in RNAlater (ThermoFisher). For each biopsy, total placental RNA was extracted from approximately 5 mg of tissue using the “mirVana miRNA Isolation Kit” (Ambion) which efficiently isolates all RNAs longer than 10 nucleotides in length, followed by DNase treatment (“DNA-free DNA Removal Kit”, Ambion). RNA quality was assessed with the Agilent Bioanalyzer and all the samples with RIN values ≥ 7.0 were used in the downstream experiments. RNA-libraries were prepared from 1µg of total placental RNA with the TruSeq Stranded mRNA Library Prep Kit (Illumina) which captures polyA-tailed transcripts by oligo-dT beads, then pooled and sequenced (single-end, 50bp) using a Single End V4 cluster kit and Illumina HiSeq2500. RNA was also extracted from human first and second trimester placental villi using the RNeasy Plus Universal Mini Kit (Qiagen). Libraries were made using the Illumina TruSeq Stranded mRNA Library Kit according to the manufacturer’s instructions.

The adaptor sequences and poor-quality bases were trimmed using *cutadapt* v1.16 (with python v3.6.1) with the following command:

*cutadapt -j 32 -a AGATCGGAAGAGCACACGTCTGAACTCCAGTCAC -q 20 -O 8 -m 20 -o $TRIMMED_FASTQ $INPUT_FASTQ*

The quality-assured trimmed short reads were mapped to the GRCh38 version of human genome reference using *TopHat2* (v2.0.12):

*tophat2 -p 32 --library-type fr-firststrand --output-dir $OUTPUT_DIR --max-multihits 10 --prefilter-multihits -transcriptome-index=$TR_INDEX $BOWTIE2INDEX $TRIMMED_FASTQ*

The transcriptome index above was built using transcript annotation from Ensembl v88 (equivalent to Gencode v26). We applied so-called two-pass (or two-scan) alignment protocol to rescue unmapped reads from the initial mapping by re-aligning unmapped reads toward the exon-intron junctions detected in the first-mapping:

*tophat2 -p 32 --library-type fr-firststrand --output-dir $OUTPUT_DIR --raw-juncs $MERGED_JUNCTION $UNMAPPED_FASTQ*

For each sample, the initial and second mapped reads were merged by *samtools* (v1.2-24-g016c62b):

*samtools merge $MERGED_BAM $FIST_MAP_BAM $SECOND_MAP_BAM*

Before gene-level quantification of read counts, we pre-processed the transcript annotation file (Gencode v26) using the ‘*collapse_annotation*.*py*’ python script available from the following GTEx github site: https://github.com/broadinstitute/gtex-pipeline/tree/master/gene_model.

*python3 collapse_annotation*.*py $GENCODE_26_GTF $PROCESSED_GENCODE_26_GTF*

Finally, we quantified sequencing reads at the gene-level using *featureCounts* tool of *subread* package (v1.5.1):

*featureCounts -T 32 -a $PROCESSED_GENCODE_26_GTF -Q 10 -s 2 -p -C -o $GENE_COUNT_OUTPUT $MERGED_BAM*

### Tissue collection for RT-qPCR analysis

Placental tissues for qPCR validation were collected from healthy women with normal term pregnancies and scheduled for delivery by elective cesarean section. Participants were consented for research sample collection as part of the surgical procedure, with further permission for storage and transfer of materials to the biobank given under approval 07/MRE05/44. Analysis was performed as part of the Cambridge Blood and Stem Cell Biobank REC ID 18/EE/0199. Human aorta, lung and left ventricle used in the research study was obtained from the Papworth Hospital Research Tissue Bank. Written consent was obtained for all tissue samples using Papworth Hospital Research Tissue Bank’s ethical approval (East of England - Cambridge East Research Ethics Committee) under approval 18/EE/0269. Human colon, stomach and small bowel were obtained from Cambridge University Hospitals Human Research Tissue Bank under approval 04/Q1604/21 and breast tissues from the Institute of Metabolic Sciences.

Approximately 35 mg of frozen tissues were homogenized by bead beating for 20 s at a speed of 4.5 ms^-1^ on a FastPrep24 sample disruption system with Lysing Matrix S tubes (MP Biomedicals, Santa Ana, CA). Total RNA was isolated with the RNeasy Plus Mini Kit (Qiagen) and 200 ng of total RNA from each sample was reverse transcribed using the High-capacity RNA-to-cDNA kit (ThermoFisher Scientific). The qPCR reactions were prepared using TaqMan Multiplex Master Mix (ThermoFisher Scientific).

### Whole genome sequencing and data processing

The whole genome sequencing dataset of the placenta (n=80) was from ‘cohort1’ (babies delivered by pre-labor Caesarean section) described in our previous report(de Goffau et al. 2019), which was also based on the POP study. Detailed description of the experimental protocol is available in the original paper.

The sequencing files were converted from CRAM format to FASTQ using *samtools* (v1.7-15-g9ce8c64):

*samtools fastq -F 0×200 $INPUT_CRAM -1 $FASTQ_R1 -2 $FASTQ_R2*

The adaptor sequences and poor-quality bases were trimmed using *cutadapt* v1.16 (with python v3.6.1) with the following command:

*cutadapt -j 32 -a AGATCGGAAGAGCACACGTCTGAACTCCAGTCAC -A AGATCGGAAGAGCGTCGTGTAGGGAAAGAGTGTAGATCTCGGTGGTCGCCGTA TCATT -q 20 -O 8 -m 20 -o $TRIMMED_R1 -p $TRIMMED_R2 $FASTQ_R1 $FASTQ_R2*

The quality-assured trimmed short reads were mapped to the GRCh38 version of human genome reference using *bwa* (v0.7.17-r1188):

*bwa mem -M -t 32 \*

*-R*

*“@RG\tID:$ID\tPL:illumina\tPU:run\tLB:$ID\tSM:$Barcode\tCN:CamObsGynae” \ $GRCh38_GENOME_FASTA $TRIMMED_R1 $TRIMMED_R2 \* | *samtools view -Sb - > $BAM_FILE*

The whole genome sequencing dataset of the 1,682 healthy normal tissues (the endometrium, blood, colon, and liver) was generated at the Wellcome Trust Sanger Institute(Moore et al. 2020).

### GTEx data processing

We compared our placenta RNA-Seq datasets with 46 somatic tissues from GTEx (v8.p2). To select eligible samples from GTEx RNA-Seq datasets, we used the same filtering conditions applied to our previous study(Gong et al. 2021): (a) RNA integrity number (SMRIN) ≥6, (b) mapping rate (SMMAPRT) ≥0.9, (c) exonic mapping rate (SMEXNCRT) ≥0.75, and (d) ≥20 qualifying samples per tissue. Five tissues (out of 54) were dropped after applying the aforementioned filters: the kidney (medulla), the fallopian tube, the cervix (endocervix), the cervix (ectocervix), and the bladder. We further removed the following three non-solid tissues: the whole blood, cultured fibroblast cells, and EBV-transformed lymphocytes cells. Finally, a total of 11,803 samples were selected from 46 somatic tissues. **Supplementary Table 2** shows the number of samples across the 46 tissues we considered. In our initial analysis, we used 4,454 samples from 20 somatic tissues from GTEx with the following modified criteria: (a) the RNA integrity number (SMRIN) □6, (b) mapping rate to genome (SMMAPRT) >0.8, (c) mapping rate to exon (SMEXNCRT) >0.8, (d) ≥10 qualifying samples of both sexes (i.e. at least 20 samples per tissue), and (e) manual selection of tissue sub-types if two or more were available for the same tissue. We considered 56,156 genes from the gene-level quantification information available from the following file: GTEx_Analysis_2017-06-05_v8_RNASeQCv1.1.9_gene_reads.gct.gz.

### Identification of absent or depleted protein-coding transcripts

We made a count matrix of 56,156 genes by 11,876 samples (i.e. 11,803 samples from GTEx and 73 from the placenta of the following gestational ages: 8 at 8wk, 6 at 14wk and 59 at term), then filtered out genes of the following conditions: (1) the sum of read count across samples is zero (n=238), (2) non-polyadenylated RNAs (e.g. transcripts of major histones) reported from the study of Yang et al.(Yang et al. 2011) (n=90), and (3) transcripts which are not annotated as ‘protein-coding’ as per Ensembl v88 (n=36,658). After filtering, a total of 19,170 protein-coding genes were considered. To adjust differences in the composition of RNA populations across multiple tissues, we applied the following two normalization methods to the count matrix (a dimension of 19,170 × 11,876): (1) the median ratio method implemented in the ‘*estimateSizeFactors*’ function of DESeq2 (v.1.26) Bioconductor package(Anders and Huber 2010), and (2) the trimmed mean of M-values (TMM), available from ‘*calcNormFactors*’ function of edgeR (v.3.28.1) Bioconductor package(Robinson and Oshlack 2010). Then, we built two matrices of normalized count per million (nCPM), each of which has a dimension of 19,170 × 11,876, using the ‘*fpm*’ and ‘*cpm*’ functions of DESeq2 and edgeR package, respectively. The columns (i.e. 11,876 samples) of the matrix were reduced to a size of 47 columns (i.e. 46 tissues from GTEx and 1 placental tissue either from early gestation or full term) by taking the mean of nCPM across samples of the same tissue. Using the placenta at term, we identified 5,632 and 5,727 genes for which the placenta was ranked 47 (i.e. bottom) based on DESeq and TMM normalization methods, respectively. Finally, we selected 762 of them (**Supplementary Table 3**) which satisfied the following three conditions: (1) nCPM >1 for the tissue ranked 46, (2) nCPM_(rank=46)_/nCPM_(rank=47)_ >3, and (3) fulfilling aforementioned conditions both in DESeq and TMM normalization methods. In the down-sampling analysis, we applied binomial sampling, using *`rbinom`* in R, to each column of the count matrix with the subsampling probability being 20 million divided by the sum of each column – a similar approach was previously introduced(Robinson and Storey 2014). This is equivalent to randomly choosing each individual mapped read with the same subsampling probability, so that the final number of down-sampled reads becomes 20 million per sample, which is the minimum number of sequencing reads in our placenta RNA-Seq dataset (**Supplementary Table 1**). Then, we applied the same approach of finding absent or depleted transcripts, as described earlier, to the down-sampled count matrix.

### Target selection for RT-qPCR validation

We selected a total of 13 transcripts to measure their abundance levels using qPCR. Eleven targets (NFIC, SNED1, GGT5, THBS2, C16orf45, ITGA7, WASF3, IGFBP6, RCAN2, SLCO3A1, and PPARGC1A) were selected by comparing our placenta RNA-Seq datasets to 20 non-placental tissues from GTEx (i.e. an initial analysis) based on the following criteria: 1) depleted in the placenta across three gestational ages (as described above), 2) fold-change (i.e. nCPM_(rank=20)_/nCPM_(rank=21; term-placenta)_) > 5, and 3) top 10 by the effect size (i.e. nCPM_(rank=20)_ -nCPM_(rank=21; term-placenta)_). PGC-1α (PPARGC1A) while it was depleted it was not within the top 10, but it was included considering its important role in regulating mitochondria. Two targets (KMO and ARMS2) were selected by comparing our placenta RNA-Seq datasets to 46 non-placental tissues from GTEx based on the following criteria: 1) annotated in MitoMiner(Smith and Robinson 2016), 2) highest nCPM in the placenta across three gestational ages (i.e. ranked within top 3), and 3) nCPM_(rank=4)_ > 1. There were 9 transcripts (KMO, ARMS2, LYPLA1, RDH13, RPIA, SLC25A35, ACSS1, NME6, and TEFM) satisfying these conditions and we selected top two by the average fold change of the term placenta compared to 46 non-placental tissues (i.e. nCPM_(term-placenta)_/nCPM_(non-placenta)_). The following 13 predesigned TaqMan assays were used: NFIC (Hs00232157_m1), SNED1 (Hs00966449_m1), GGT5 (Hs00897715_m1), THBS2 (Hs01568063_m1), C16orf45 (Hs01014981_m1), ITGA7 (Hs01056475_m1), WASF3 (Hs00903488_m1), IGFBP6 (Hs00181853_m1), RCAN2 (Hs00195165_m1), SLCO3A1 (Hs00203184_m1), PPARGC1A (Hs00173304_m1), ARMS2 (Hs01394203_m1), and KMO (Hs00175738_m1). The above target genes were normalized to the geometric mean of CDC34 (Hs00362082_m1) and TBP (Hs00427620_m1).

### Calculation of mitochondrial copy number

The mitochondrial copy number (*Copy*_*mt*_) was calculated as the ratio of mitochondrial depth of coverage (*Cov*_*mt*_) over the average genome depth of coverage (*Cov*_*g*_). It is formally defined as following:

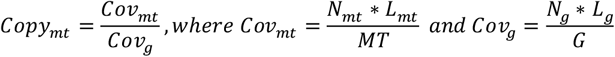

The depth of coverage (*Cov*) above is defined as the number of mapped bases, which is the number of reads (*N*) multiplied with the length of the mapped reads (*L*), divided by the haploid size of genome (*G*) or mitochondrion (*MT*). We calculated the depth of coverage from the BAM files of whole genome sequencing datasets using *bedtool* with the following command:

*bedtools genomecov -ibam $BAM_FILE > ${BAM_FILE%*.*bam}*.*cov*.*txt*.

### Gene Ontology analysis

The gene ontology analysis was performed using g:Profiler(Raudvere et al. 2019) (https://biit.cs.ut.ee/gprofiler; version e103_eg50_p15_68c0e33) with FDR multiple-testing correction method applying significance threshold of 0.05. We used the *gprofiler2* R package (https://cran.r-project.org/web/packages/gprofiler2/), a R client for the g:Profiler tools, with the 19,170 protein-coding genes (see above) as a list of background genes.

### Dimension reduction

We used *Rtsne* (https://cran.r-project.org/web/packages/Rtsne; v0.15) with a default option for the main dimension reduction method as shown in **Figure 3C**.

### External datasets used in this study

#### mtDNA copy numbers in cancer

we downloaded mtDNA copy numbers of the Cancer Genome Atlas Pan-Cancer Analysis of Whole Genomes (PCAWG) Consortium(Yuan et al. 2020) from the following URL: https://ibl.mdanderson.org/tcma/download/TCMA-CopyNumber.tsv.zip. The dataset was downloaded as of the 9^th^ of March 2020.

#### WGS datasets of healthy human tissues

we used WGS datasets of healthy human tissues from the European Genome-phenome Archive (EGA, https://ega-archive.org/) with the following accession numbers: EGAD00001004086 (blood), EGAD00001004192 (colon), EGAD00001004547 (endometrium), and EGAD00001004578 (liver).

#### Placental RNA-Seq datasets

we downloaded four placenta RNA-Seq datasets from the European Nucleotide Archive (ENA, https://www.ebi.ac.uk/ena) with the following accession numbers: PRJNA386110 (Lim), PRJNA499121 (Huang), PRJNA704615 (Ashley), and PRJNA472249 (Awamleh). The RNA-Seq dataset from the Verheecke’s study was obtained personally from one of the authors.

### Identification of transcripts localized in the mitochondria

We downloaded MitoMiner (v4), a dataset of mitochondrial localization, from the following URL: http://mitominer.mrc-mbu.cam.ac.uk/release-4.0/mitocarta.do. We selected genes of the following conditions: (1) not encoded in mitochondrial chromosome, (2) “Known mitochondrial” or “Predicted mitochondrial” as types of evidence, and (3) one of the 19,156 eligible protein-coding genes described above. After filtering, we considered 1,042 protein-coding genes.

### Code Availability

Codes used in this study is available in the Methods section and at https://gitlab.com/sunggong/pops-placenta-mt-2020.

## Data access

The term placenta RNA-Seq data have been deposited in the European Genome-phenome Archive (EGA, https://ega-archive.org/) with the following accession number: EGAD00001006304. The early gestation placenta RNA-seq data have been deposited in the European Nucleotide Archive (ENA, https://www.ebi.ac.uk/ena) with the following accession number: PRJEB38810. The term placenta WGS data have been deposited in the EGA with the following accession number: EGAD00001004198. Correspondence and requests for materials should be addressed to D.S.C-J.

## Acknowledgements

This work was supported by the Medical Research Council (United Kingdom; G1100221 and MR/K021133/1) and the National Institute for Health Research (NIHR) Cambridge Biomedical Research Centre (Women’s Health theme). I.A. is funded by the Centre for Trophoblast Research (CTR) Next Generation Fellowship. G.A. is funded by the CTR PhD scholarship. A.R.J.L. and L.M.R.H. are funded by Wellcome PhD studentships. We would like to thank Katrina Holmes, Josephine Gill, Leah Bibby, Samudra Ranawaka and Ryan Millar for technical assistance during the study. We would like to thank Dr Álvaro Cortés-Calabuig for kindly sharing research data. The views expressed are those of the authors and not necessarily those of the NHS, the NIHR or the Department of Health and Social Care.

## Author contributions

D.S.C-J. and G.C.S.S. conceived the experiments. S.G, D.S.C-J, G.C.S.S, designed the experiments. F.G, I.A, G.A. and E.C. performed the experiments. S.G. analyzed all the sequencing data. E.C. managed sample collection and processing and the biobank in which all samples were stored. A.R.J.L. and L.M.R.H. provided and processed sequence data. All authors contributed to writing the manuscript and approved the final version.

